# Timing of starvation determines its effects on susceptibility to infection in *Drosophila melanogaster* females independent of host evolutionary history

**DOI:** 10.1101/2024.09.11.612402

**Authors:** Aabeer Basu, Aparajita Singh, Nagaraj Guru Prasad

## Abstract

An organism’s susceptibility to pathogens is contingent on various environmental factors, including the availability of nutrition. Starvation can alter host susceptibility to infections, either directly via depletion of resources essential for proper functioning of the immune system, or indirectly via the various physiological changes it induces within the host body. We tested if the susceptibility of *Drosophila melanogaster* populations to *Enterococcus faecalis* infection is affected by (a) whether the hosts are starved before or after the infection, and (b) the evolutionary history of the host. Hosts from laboratory fly populations that have been experimentally evolved to be more resistant to *E. faecalis*, and their corresponding control populations, were subjected to infection with or without being starved prior to and after being infected. We found that the effect of starvation on susceptibility to *E. faecalis* changed with the timing of starvation: starvation after infection improved survival of infected hosts, irrespective of how they were treated before infection, while starving only prior to infection (and not after) compromised post-infection survival. The changes in infection susceptibility were uniform in both the evolved and the control populations, suggesting that the effects of starvation are not dependent on pre-existing resistance to the infecting pathogen.

## 1. Introduction

The host immune function and the expression of costs associated with it are both contingent on the availability and accessibility of nutrition and resources to the host (Sandland and Minchella 2003, Lazzaro and Little 2008). Lack of nutrition, and therefore energy and resources – either via alteration of the host environment, or due to manipulation or dysfunction of the host physiology – can compromise the host’s ability to defend against invading pathogens and increase the cost of this defense in the form of trade-offs with other physiological functions (Moret and Schmid-Hempel 2000, McKean et al., 2008, Ayres and Schneider 2009, Bashir-Tanoli and Tinsley 2014, Buchanan et al., 2018). Manipulation of host diet, both qualitative and quantitative, and thereafter studying its effects on host immune function is a tractable experimental set-up in insect model systems that have yielded significant understanding regarding the role of resource availability in affecting host immune capabilities (Ponton et al., 2013, Cotter and Al Shareefi 2022, Ponton et al., 2023).

Starvation (i.e., complete unavailability and inaccessibility of nutrition) in insects is known to alter the immune function of the host, either directly via altering the amount of resources available to the host or indirectly via alterations of the host physiology, and therefore change host susceptibility to infections (Adamo 2021). Starvation commonly depletes bodily resources and reduces the efficiency of the host immune system in the insects, thereby making the host more susceptible to pathogenic infections (Brown et al., 2000, Siva-Jothy and Thompson 2002, Banville et al., 2012, Adamo et al., 2016). The effect of starvation on the host immune system is not uniform across all its constituent components which can lead to pathogen specific effects (Adamo et al., 2016).

There is limited understanding of how starvation impacts host disease susceptibility and what other factors can modify the effects of starvation in *Drosophila melanogaster*. Starvation in immune-deficient flies has been shown to improve host post-infection survival for Gram-negative bacterial pathogens (but not Gram-positive bacteria; Brown et al., 2009), whereas starvation in wild-type flies can have variable effects on post-infection survival of the host contingent upon both the identity of the infecting pathogen and the host mating status (Basu et al., 2022). These studies also differed in terms of the timing of starvation: the former study starved the hosts for 24 hours prior to infection while the later study starved the hosts from the point of infection onwards (Brown et al., 2009, Basu et al., 2022). At the sub-organismal level, starvation has been reported to down-regulate immune genes (Harbison et al., 2005), increase production of antimicrobial peptides (Becker et al., 2010), and both reduce (Brown et al., 2009) and increase (Basu et al., 2022) proliferation of bacterial pathogens within host body, suggesting mixed results across individual studies in *D. melanogaster*. Overall, it is difficult to predict the effect of starvation on susceptibility to any specific pathogen (Adamo 2021).

In this study, using laboratory populations of *D. melanogaster*, we tested if the effect of starvation on host susceptibility to pathogenic bacterial infection is influenced by (a) the timing of starvation (before or after) relative to the point of infection, and (b) the evolutionary history of the host, that is, when the host is adapted to pathogenic infections. Dietary manipulations have been shown to exert different effects on host immune function depending upon if the manipulation was implemented before or after infection in *D. melanogaster* (McKean and Nunney 2005). The effect of dietary manipulations on *D. melanogaster* immune function is also known to be dependent on host genetic make-up (Howick and Lazzaro 2014, Unckless et al., 2015). And, adaptation to pathogenic infections can alter how hosts respond to phenotypic manipulations that can affect their susceptibility to infections (Basu et al., 2024a).

We experimentally evolved replicate laboratory populations of *D. melanogaster*, selecting for flies that survive pathogenic infection with the Gram-positive bacterium *Enterococcus faecalis*. The selected populations evolved to have greater post-infection survival, and greater capacity to limit within-host proliferation of the infecting bacteria compared to the control populations (Singh et al., 2021, Basu et al., 2024b). Using females from these selected populations and their ancestrally paired control populations (**see section 2.1**) we investigated if host adaptation to pathogenic infection alters the effects of starvation on host susceptibility to infection with the native pathogen (i.e., the pathogen used during selection, *E. faecalis*). Our results demonstrate that starvation alters the susceptibility of female flies to *E. faecalis* infection, contingent on whether the flies were subjected to starvation before or after infection, but the effects of starvation are not dependent on whether the host is adapted to the infecting pathogen.

## 2. Materials and methods

### 2.1. EPN experimental evolution set-up

The experiments reported in this study were carried out using flies from the EPN experimental evolution set-up consisting of twelve populations distributed across three selection regimes (previously described in Singh et al., 2021).

#### E regime

The E_1-4_ populations are selected for better survival following infection with the Gram-positive bacterium *Enterococcus faecalis*. 2–3-days old adult flies (200 females and 200 males) are subjected to infection with *E. faecalis* every generation, and 96-hours post-infection, the survivors are allowed to reproduce and contribute towards the next generation. At the end of 96 hours, on average 100 females and 100 males are left alive in each of the E_1-4_ populations.

#### P regime

The P_1-4_ populations are procedural (sham-infected) control populations. 2–3-days old adult flies (100 females and 100 males) are subjected to sham-infections every generation, and 96-hours post-sham-infection, the survivors are allowed to reproduce and contribute towards the next generation.

#### N regime

The N_1-4_ populations are uninfected control populations. 2–3-days old adult flies (100 females and 100 males) are subjected only to light CO_2_ anesthesia every generation, and 96-hours post-handling, the survivors are allowed to reproduce and contribute towards the next generation. (Under usual circumstances, negligible mortality occurs in the P_1-4_ (< 2%) and N_1-4_ (0%) populations during maintenance of the selection regimes.) These populations were derived from the ancestral Blue Ridge Baseline (BRB_1-4_) populations (creation and maintenance of the BRB populations are described in Singh et al., 2015). The E_1_, P_1_, and N_1_ populations were derived from the BRB_1_ population and constitute ‘block 1’ of the experimental evolution regime. Similarly, E_2_, P_2_, and N_2_ populations were derived from the BRB_2_ population and constitute ‘block 2’, and so on. This block design implies that E_1_, P_1_, and N_1_ have a more recent common ancestor, compared to E_1_ and E_2_, or P_1_ and P_2_, and so on. Populations belonging to each block were handled together, both during maintenance of the populations and during experiments. Please refer to **supplementary figure 1** for a graphical depiction of the block design.

The regular maintenance of the EPN set-up has been previously described (Singh et al., 2021, Basu et al., 2024b). The populations are maintained on a diet of banana-jaggery-yeast food medium. Every generation, eggs are collected at a density of 60-80 eggs per vial, in 6-8 ml food medium. 10 such rearing vials (9 cm height × 2.5 cm diameter) are set up for each of the 12 populations. These vials are incubated at 25 ^O^C, 60% RH, and a 12:12 LD cycle. Under these conditions, eggs develop into adults within 9-10 days of egg collection. The adults stay in the rearing vial till day 12 post-egg laying (PEL); by this point of time the flies are sexually mature and have already mated. On day 12 PEL, flies from each population are handled according to their regime, as described above (**see section 2.2 for details of the infection process**). The flies are thereafter housed in plexiglass cages (14 cm × 16 cm × 13 cm); one cage for each population. The cages are provided with fresh food medium, on a 60 mm Petri plate, on every alternate day. On day 16 PEL the surviving flies are allowed to lay eggs for the next 18 hours on a fresh food plate in each cage. Eggs are collected from these oviposition plates to start the next generation.

### 2.2. Pathogen handling and infection protocol

*Enterococcus faecalis* (**Lazzaro et al., 2006**), a Gram-positive bacterium, was used in regular maintenance of the EPN populations and during the experiment. During regular maintenance of the EPN populations, the flies from the E_1-4_ populations (**see section 2.1**) are infected with an *E. faecalis* suspension of OD_600_ = 1.5. For all experimental infections (**see section 2.4**), flies were also infected with an E*. faecalis* suspension of OD_600_ = 1.5 to keep experimental conditions consistent with that of regular maintenance. OD_600_ = 1.0 corresponds to 10^6^ cells/ml for *E. faecalis*.

The bacteria are stored as glycerol stocks (17%) in -80 ^O^C. To obtain live bacterial cells for infections, 10 ml lysogeny broth (Luria Bertani Broth, Miler, HiMedia) is inoculated with glycerol stocks and incubated overnight with aeration (150 rpm shaker incubator) at 37 ^O^C. 100 microliters from this primary culture is inoculated into 10 ml fresh lysogeny broth and incubated for the necessary amount of time to obtain confluent (OD_600_ = 1.0-1.2) cultures. The bacterial cells are pelleted down using centrifugation and resuspended in sterile MgSO_4_ (10 mM) buffer to obtain the required optical density (OD_600_) for infection. Flies are infected, under light CO_2_ anaesthesia, by pricking them on the dorsolateral side of their thorax with a 0.1 mm Minutien pin (Fine Scientific Tools, USA) dipped in the bacterial suspension. Sham-infections (injury controls) are carried out in the same fashion, except by dipping the pins in sterile MgSO_4_ (10 mM) buffer.

### 2.3. Pre-experiment standardization and generation of experimental flies

Prior to experiments, flies from the different selection regimes were reared for one generation under ancestral maintenance conditions to account for potential non-genetic parental effects (Rose 1984); flies thus generated are referred to as *standardized* flies. To generate standardized flies, eggs were collected from all populations at a density of 60-80 eggs per vial (with 6-8 ml food medium); 10 such vials were set up per population. The vials were incubated under standard maintenance conditions (**section 2.1**). On day 12 PEL, the adults were transferred to plexiglass cages (14 cm × 16 cm × 13 cm) with food plates (Petri plates, 60 mm diameter). Eggs for experimental flies were collected from these *standardized* population cages.

To generate flies for the experiments, eggs were collected from *standardized* population cages at a density of 60-80 eggs per vial (with 6-8 ml food medium); 60 vials were collected per regime per block. The vials were incubated under standard maintenance conditions. The eggs developed into adults by day 10 PEL. The adults continued to be housed in these vials till day 12 PEL when they were distributed into respective experimental treatments (**section 2.4**). The handling of the flies during the course of the experiment was kept identical to how the populations are regularly maintained to keep their ecology consistent.

### 2.4. Experiment design

Only flies from N and E selection regimes were used in the experiment reported here. Previous works have reproducibly demonstrated that N (uninfected control regime) and P (procedural control regime) flies do not differ significantly in terms of post-infection survival (Singh et al., 2021, Singh et al., 2022a, Singh et al., 2022b, Basu et al., 2024b). This experiment was carried out after 80 generations of forward selection.

On day 12 PEL, 2–3-day old flies flies from N and E regimes were randomly assigned to one of the four following treatments:

a. *Fed/Fed* (FF): flies that were continuously fed their standard diet.
b. *Fed/Starved* (FS): flies that were fed their standard diet prior to infection, and subjected to starvation post infection.
c. *Starved/Fed* (SF): flies that were subjected to starvation for 48 hours prior to infection, but fed their standard diet post infection.
d. *Starved/Starved* (SS): flies that were continuously starved starting from 48 hours prior to infection.

15 vials worth of flies were allocated for each selection regime (N and E) × starvation treatment (FF, FS, SF, and SS) × block (1, 2, 3, and 4) combination. Flies in the FF and FS treatments were transferred into fresh food vials (with 2 ml standard food medium) from their rearing vials on day 12 PEL. Flies in the SF and SS treatments were transferred into *starvation* vials (with 2 ml non-nutritive 2% agar gel) from their rearing vials on day 12 PEL. Flies were held in these vials till the time of infection which was 48-50 hours after this transfer.

Flies were subjected to infection or sham-infection on day 14 PEL when they were 4-5 days old as adults. Flies in each vial were anesthetized with CO_2_, and 8 females were infected (10 vials) or sham-infected (5 vials) from each vial (**as described in section 2.2**), and placed in a fresh vial after handling. Females in the FF and SF treatments were placed in fresh food vials while females in the FS and SS treatments were placed in fresh *starvation* vials. The females were housed in these vials for the next 48 hours while their mortality was recorded every 4 to 6 hours. In total, 80 females (per selection regime per starvation treatment per block) were infected while 40 females (per selection regime per starvation treatment per block) were sham-infected.

Experiments for each block were carried out on a separate day.

### 2.5. Statistical analysis

Data analysis was carried out using R statistical software (version 4.1.0; R Core Team 2021). Survival data of infected flies was modeled as

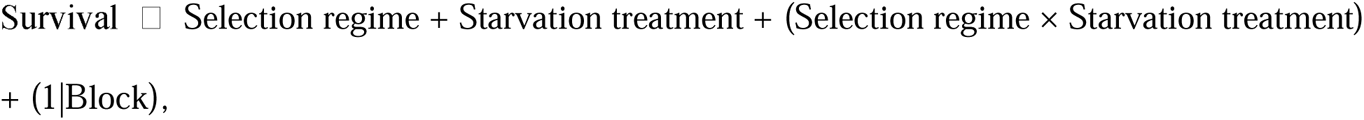

using mixed-effects Cox proportional hazards (*coxme* function from *coxme* package; Therneau 2020), where selection regime, starvation treatment, and their interaction were modeled as fixed factors, and block identity was modeled as a random factor. This model was subjected to analysis of deviance (type II; *Anova* function from *car* package; Fox and Weisberg 2019) for significance testing for the fixed factors. Results for the analysis of deviance are reported in **table 1** and the output of the Cox proportional hazards model is reported in **supplementary table 1**. Since there was negligible mortality in sham-infected flies in both selection regimes and in all starvation treatments (**figure 1**), only the survival data of infected flies were subjected to statistical analysis.

**Figure 1.**
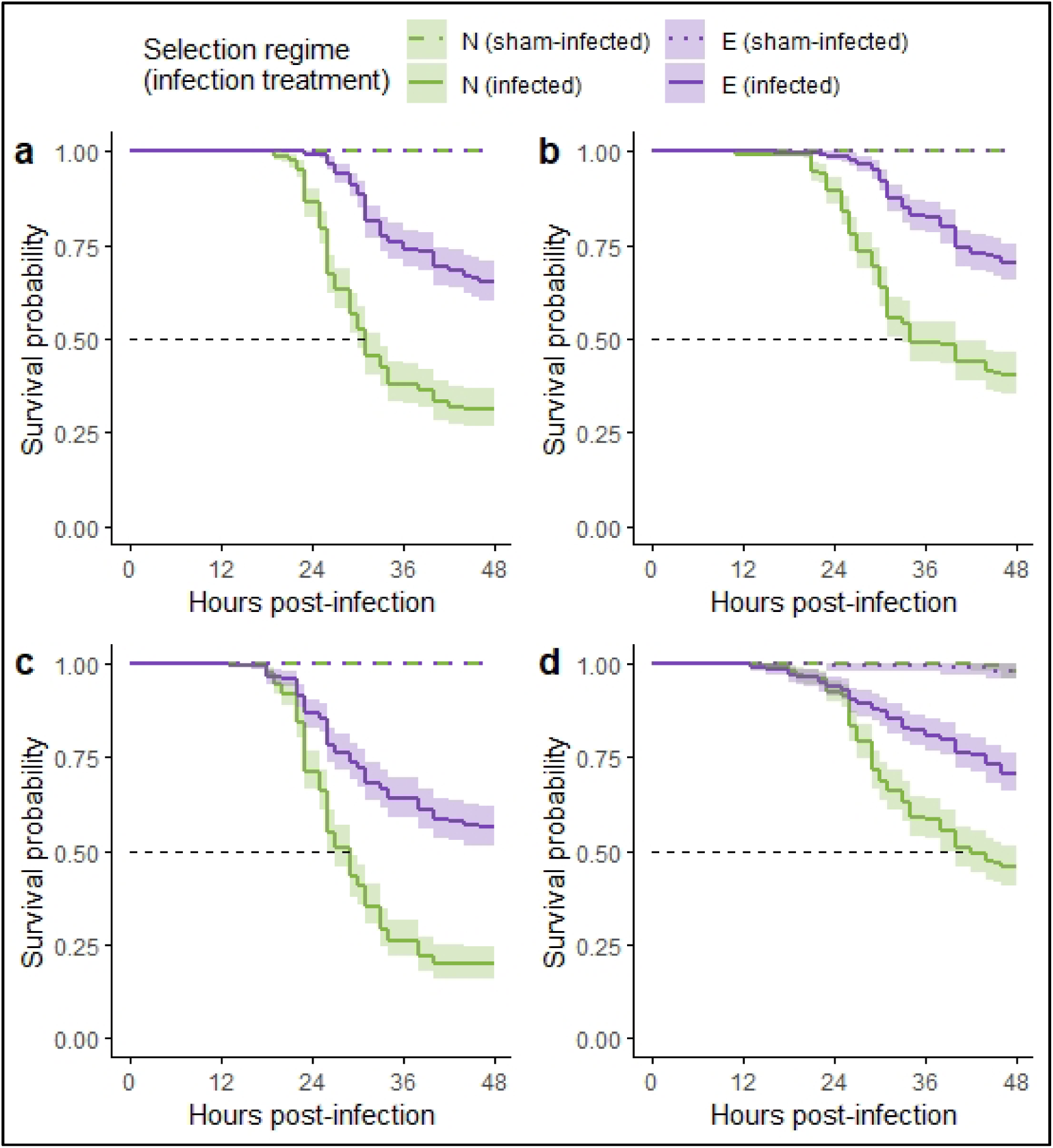
Effect of selection history (regime) and starvation treatments on survival of *Drosophila melanogaster* females from the N (control) and E (selected) regimes when infected with *Enterococcus faecalis* or sham infected. **(a)** *Fed/Fed*; **(b)** *Fed/Starved*; **(c)** *Starved/Fed*; and **(d)** *Starved/Starved*. (Survival curves plotted using Kaplan-Meier method after pooling data across all four blocks for each selection regime. The shaded region represents a 95% confidence interval around the survival curves. The horizontal dashed line in individual panels demarcates 50% survival probability.)

**Table 1.** Analysis of deviance (type II) for the effect of selection regime, starvation treatment, and their interaction on survival of *Drosophila melanogaster* females infected with *Enterococcus faecalis*.

| Fixed factors | DF | Chi-square value | p-value |
| --- | --- | --- | --- |
| Selection regime | 1 | 301.2692 | < 0.0001 |
| Starvation treatment | 3 | 117.3288 | < 0.001 |
| Selection regime × Starvation treatment | 3 | 3.9737 | 0.2643 |

## 3. Results

Selection regime (p < 0.001) had a significant effect on post-infection survival of females (**figure 1**, **table 1**): E regime females had an overall lower risk of mortality relative to N regime females (hazard ratio, 95% confidence interval: 0.3291, 0.2618 – 0.4136). Starvation treatment (p < 0.001) also had a significant effect on survival of infected females (**table 1**): *Fed/Starved* and *Starved/Starved* females had lower risk of mortality relative to *Fed/Fed* females, while Starved/Fed females had greater risk of mortality (**figure 2, supplementary table 1**). Selection regime x starvation treatment interaction did not have a significant effect on post-infection survival (**table 1**).

**Figure 2.**
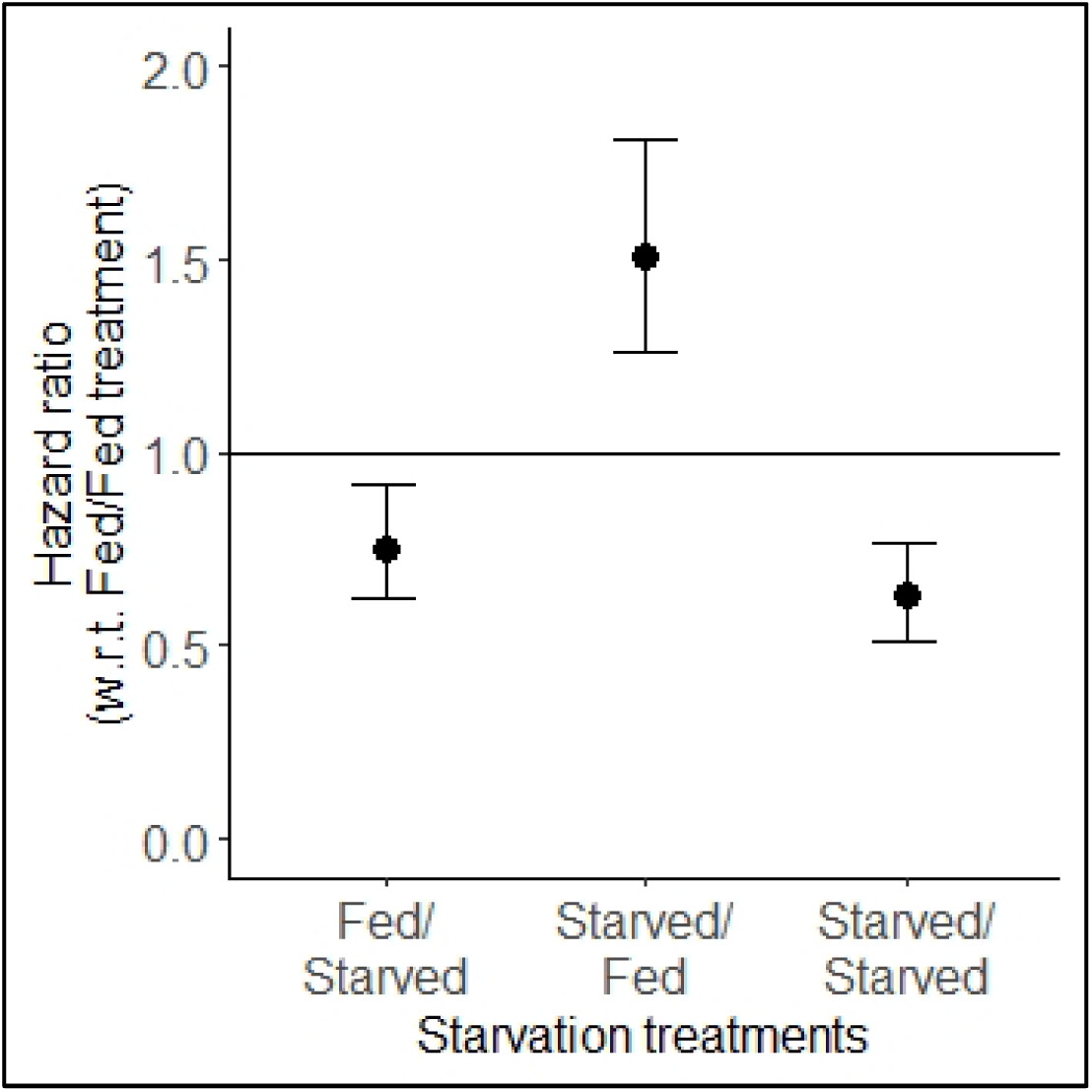
Effect of starvation on mortality of *Drosophila melanogaster* females infected with *Enterococcus faecalis*. Hazard ratio (mixed-effects Cox proportional hazards model) for females from different starvation treatments, relative to females from *Fed/Fed* treatment, across both selection regimes. The hazard for *Fed/Fed* treatment is normalized to 1, and represented by the horizontal solid line on the plot. (Error bars represent 95% confidence intervals.)

## 4. Discussion

In this study we explored the differential effects of starvation on susceptibility to *Enterococcus faecalis* infection in *Drosophila melanogaster* populations experimentally evolved for increased survival following *E. faecalis* infection. The effect of dietary manipulations on host immune function in *D. melanogaster* flies can potentially be determined by whether the manipulation was implemented before or after the infection (McKean and Nunney 2005). In our experiment, therefore, the hosts were either (a) continuously fed, (b) starved only after the infection, (c) starved only before the infection but not after, or (d) continuously starved both before and after the infection. Females from the selected (E_1-4_ populations) and the uninfected control (N_1-4_ populations) regimes were subjected to these four treatments and their post-infection survival was quantified.

Previous work with these populations has demonstrated that malnutrition has selection regime specific effects on female post-infection survival: malnutrition does not have an effect on post-infection survival of females from the control regimes, but counterintuitively improves post-infection survival of females from the selected regime (Singh et al., 2022a). Females from these selection regimes, additionally, do not differ in terms of their baseline starvation resistance (Singh et al., 2022b). The present work builds on the existing work to test if complete deprivation of nutrition, in the form of starvation, has any differential effect on post-infection survival in these populations. Although both starvation and malnutrition represent deprivation of nutrition, their effects on host immunity are not always identical in insects (Adamo et al., 2016).

### 4.1. Effect of starvation depends on the timing of starvation relative to infection

Our *a priori* expectation was that the post-infection survival of the females would be compromised by starvation, and the effect would increase with increasing duration of starvation. Starvation is known to compromise immune function in a variety of insects for both juveniles and adults (viz., Brown et al., 2000, Siva-Jothy and Thompson 2002, Banville et al., 2012, Adamo et al., 2016). In the case of *D. melanogaster*, previous studies have reported that the effect of starvation on survival following bacterial infection can be pathogen specific, and contingent on other host factors (viz. mating status) (Brown et al., 2009, Basu et al., 2022).

We observe in our experiments that the effect of starvation on post-infection survival depends on whether starvation was implemented before or after infection. In general, starving the females after the infection improved post-infection survival irrespective of how they were handled prior to infection. There was a 24% increase in post-infection survival, relative to the continuously fed females (*Fed/Fed treatment*), when females were fed before infection but starved afterwards (*Fed/Starved treatment*), while continuously starving the females both before and after infection (*Starved/Starved treatment*) lead to a 37% increase in post-infection survival (**figures 1.b, 1.d, and 2**). Increased survival due to starvation after infection agrees with previous reports for *D. melanogaster* females infected with *E. faecalis*, that post-infection starvation boosts survival when the females are mated (Basu et al., 2022); the females in our experiments were mated by the time of infection. But greater increase in post-infection survival in *Starved/Starved* treatment is unexpected and counter-intuitive since females in this treatment had been starved to the greatest extent (96 hours by the end of the experiment). We know of no previous studies that have demonstrated that the survival of *E. faecalis* infected flies improves due to quantitative resource limitation of the host in any form, except in a previous experiment with these same populations (Singh et al., 2022a).

Starving the females only before the infection (*Starved/Fed treatment*) increased post-infection mortality of the females by about 51% compared to the continuously fed (*Fed/Fed* treatment) females (**figures 1.c and 2**). Importantly, our results thus disagree with what has been reported by Brown and colleagues (2009) who starved flies briefly before infection and found that susceptibility to *E. faecalis* infection remains unaffected. It is noteworthy that the previous study used immune-deficient inbred hosts while our study involves wild-type outbred hosts. Our results agree with observations in beetles (Siva-Jothy and Thompson 2002) where the effects of starvation on host immune function can be erased, or in our case reversed, when hosts are allowed to feed again after being starved for a brief period of time.

Overall, although previous studies have demonstrated that nutrition limitation can improve survival of *D. melanogaster* flies after infection with certain pathogens (viz. Burger et al., 2007, Ayres and Schneider 2009), we report here for the first time that such improvement is contingent upon the timing of resource limitation relative to infection.

Starvation had negligible effect on survival of sham-infected females during the course of our experiment (**figure 1**), with mortality (albeit minimal) being observed only in the *Starved/Starved treatment* females (**figure 1.d**). This was not surprising since previous work has shown that flies of these populations start dying of starvation only after being starved for at least 5 to 6 days (Singh et al., 2022b). Hosts in our experiments were starved for a maximum of four days by the end of the experiment (in the *Starved/Starved treatment*); in other treatments, the hosts were starved for even less time. Hence, an absence of effect of starvation on survival of sham-infected females agrees with what was known previously about these populations. No death in sham-infected females additionally suggests that we are within the time range where death due to starvation has not yet started to occur and all recorded deaths are likely to be due to infection.

### 4.2. Effect of starvation is independent of host evolutionary history

We originally expected that the effect of starvation on post-infection survival would depend on the host evolutionary history. Hosts adapted to survive pathogenic infections are expected to have evolved to invest extra resources towards disease resistance, and the cost of this excess investment becomes apparent when resources are scarce (Boots 2011, Fellowes et al., 1998, Faria et al., 2015). Adapted hosts can therefore be expected to suffer greater loss of fitness (post-infection survival) due to starvation. Alternatively, if the adapted hosts have evolved to optimize their immune function and buffer against environmental fluctuations that compromise immunity, adapted hosts are expected to be affected by starvation to a lesser degree compared to non-adapted hosts (Basu et al., 2024a).

Contrary to expectations we did not detect any ‘selection regime × starvation treatment’ interaction in our experimental data. This indicates that the effects of starvation treatments did not differ between the E regime and the N regime females. Our results therefore suggest that the effect of starvation on host susceptibility to *E. faecalis* infection is not modified by host adaptation, and therefore greater resistance, to pathogenic infection by the same bacteria. This is surprising since previous studies have reported that the effect of dietary manipulations on host immune function can depend on the host genotype in *D. melanogaster* (McKean et al., 2008, Howick and Lazzaro 2014, Unckless et al., 2015). One important difference between our study and the previous ones is that the previous studies either manipulated the levels of protein (McKean et al., 2008) or carbohydrate (Howick and Lazzaro 2014, Unckless et al., 2015) in the food medium while we starved our flies, subjecting them to complete resource deprivation.

### 4.3. Speculations, limitations, and future directions

Given the observations from our experiment, that (a) prolonged starvation does not compromise post-infection survival and (b) feeding after being starved does compromise survival, we speculate that the effect of starvation on susceptibility to *E. faecalis* infection is not determined by the effect of starvation on host resource stores. Rather the effect may be driven by various physiological changes and metabolic signaling that are altered due to starvation within the host (Adamo 2021, Ponton et al., 2023). For example, starvation suppresses egg production in *D. melanogaster* females (Bownes et al., 1988, Terashima and Bownes 2004), which can free up resources that can be re-routed towards immune defense in starved females, while fed females continue to allocate resources towards egg production. Another possibility is that defense against *E. faecalis* was boosted by starvation during the course of infection since defense against *E. faecalis* infection is primarily mediated by constitutive defenses (Nehme et al., 2011), which have been hypothesized to be prioritized during resource limitation (Adamo et al., 2016, Adamo 2017).

One caveat of our experimental design is that *E. faecalis* seems to be a unique pathogen: various studies have reported that survival of *E. faecalis* infected flies are relatively less affected by various types of dietary manipulations, such as reduced larval nutrition (Ayres and Schneider 2009, Singh et al., 2022b), adult starvation (Brown et al., 2009, Basu et al., 2022), adult dietary restriction (Libert et al., 2008), and increased sugar content in adult diet (Darby et al., 2024), compared to survival of flies infected with other known entomopathogenic bacteria. But since *E. faecalis* was the pathogen used in experimental evolution of the E regime populations, we used the same pathogen for our experiments here, to keep the experiments ecologically consistent with the maintenance regime of our populations. Future studies must explore the effect of starvation on post-infection survival in similar experimentally evolved populations that were selected with different pathogens (viz. Ye et al., 2009, Faria et al., 2015, Gupta et al., 2016, Ahlawat et al., 2022) to confirm the generalizability of our results across different pathogens and populations.

A second caveat of our experimental design is that the females were subjected to starvation for arguably very brief periods of time: 48 hours for *Fed/Starved* and *Starved/Fed* treatments and 96 hours for *Starved/Starved* treatment by the end of the experiment. Prolonging the duration of starvation might produce different results from what we have observed: acute and chronic starvation can impact the host physiology, and therefore their immune function, differently (Rion and Kawecki 2007, Zhang et al., 2019). Prolonging the period of starvation post infection is difficult since after a point of time death due to starvation would start to confound death due to infection. This is also why we limited our experiments to 48 hours post-infection. (Additionally, 0 to 48 hours post-infection is the acute phase of *E. faecalis* infection in our study populations under usual conditions (Basu et al., 2024b).) The period of starvation before infection can be prolonged, but that would imply infecting older flies; susceptibility to infections is known to change with age in *D. melanogaster* (Corbally and Regan 2022). A delayed infection in our experiment would also have implied infecting the flies at a much later date compared to when the E (selected) regime flies are infected during their regular maintenance.

### 4.4. Conclusion

Previous research has documented that resource limitation can be either beneficial or detrimental for disease resistance in a pathogen specific manner in insects and other organisms (Hite et al., 2020). In *Drosophila melanogaster* too, such pathogen-specific effects are known for various dietary manipulations (viz. Ayres and Schneider 2009, Libert et al., 2008), including starvation (Basu et al., 2022). We have demonstrated here that resource limitation in the form of starvation can make *D. melanogaster* flies more, or less, susceptible to the same bacterial pathogen depending upon if the flies were subjected to starvation before or after infection. Starvation during the course of the infection promotes survival, irrespective of the pre-infection starvation. On the other hand, starvation prior to infection, but not during, compromises survival. Additionally, the effect of starvation on host disease resistance is not altered based on if the host is more resistant to the infecting pathogen. Our results allude to the possibility that the effect of starvation on susceptibility to bacterial pathogens in flies may not be a linear consequence of resource depletion, and might be a product of complex physiological changes that a fly undergoes when subjected to starvation.

---

**Supplementary figure 1.**
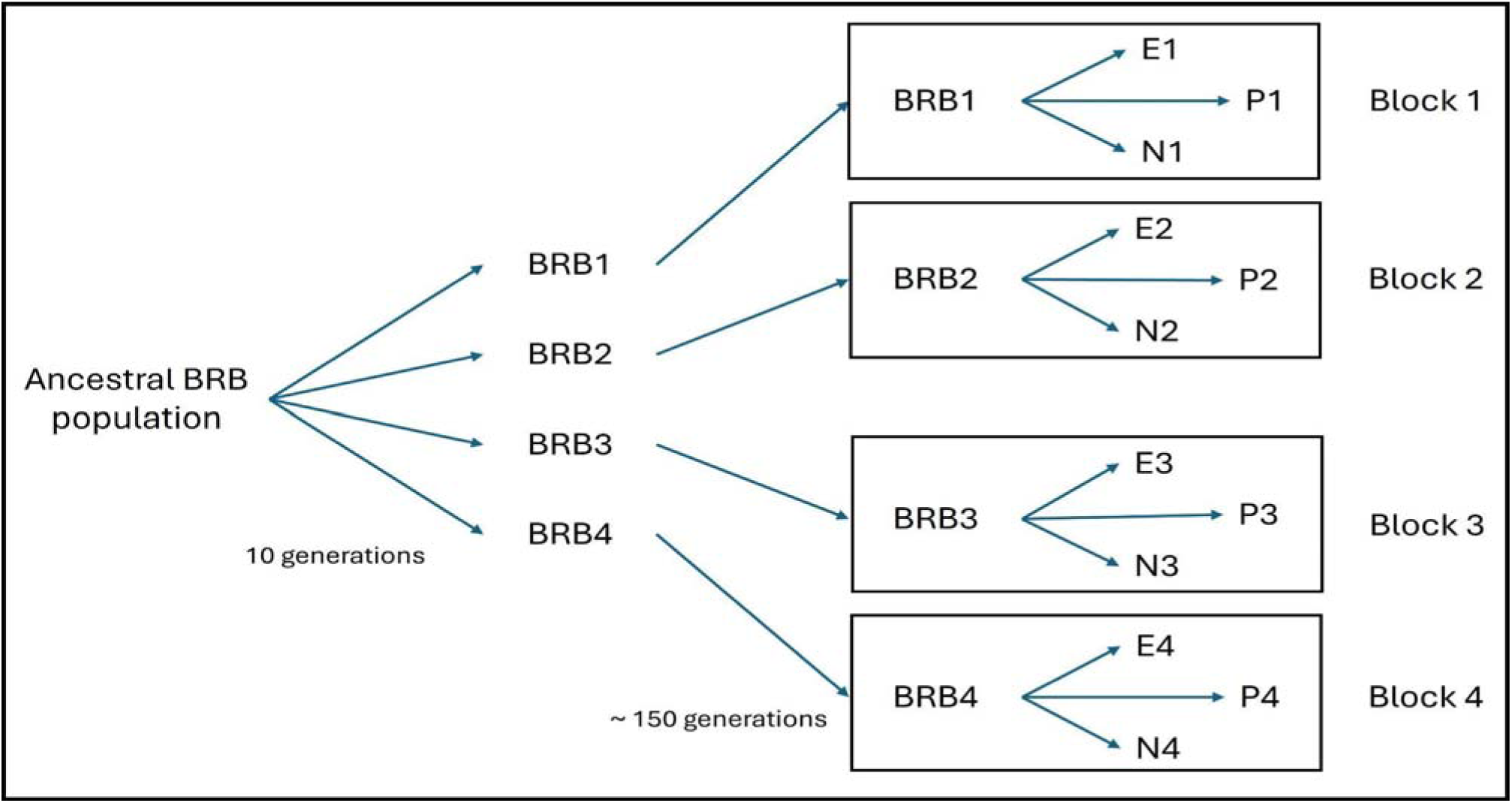
The schematics of the experimental evolution set-up depicting the block design, and the interrelatedness of the populations used in this study. Reproduced from Basu et al., 2024 (Zoology).

**Supplementary table 1.** Output of mixed-effects Cox proportional hazards analysis for females of N and E regimes subjected to various starvation treatments and infected with *Enterococcus faecalis*. Hazard ratios (HR) are relative to the default level for each factor, which is set at 1. The default level for “Selection regime” is ‘N’ and the default level for “Starvation treatment” is ‘Fed/Fed’. Hazard ratio greater than 1 implies reduced survival compared to the default level and vice versa.

| **Fixed factors (level)** | **HR** | **Lower CI** | **Upper CI** | **Z value** | **p-value** | **Variance for random factor (Block)** |
| --- | --- | --- | --- | --- | --- | --- |
| *Selection regime* (E) | 0.3291 | 0.2618 | 0.4136 | -9.53 | < 0.001 | 0.0318 |
| *Starvation treatment* (Fed/Starved) | 0.7533 | 0.6205 | 0.9146 | -2.86 | 0.004 |  |
| *Starvation treatment* (Starved/Fed) | 1.5118 | 1.2608 | 1.8128 | 4.46 | < 0.001 |  |
| *Starvation treatment* (Starved/Starved) | 0.6288 | 0.5151 | 0.7675 | -4.56 | < 0.001 |  |
| *Selection regime* (E) : *Starvation treatment* (Fed/Starved) | 1.0891 | 0.7787 | 1.5238 | 0.50 | 0.620 |  |
| *Selection regime* (E) : *Starvation treatment* (Starved/Fed) | 0.9960 | 0.7321 | 1.3552 | -0.03 | 0.980 |  |
| *Selection regime* (E) : *Starvation treatment* (Starved/Starved) | 1.3469 | 0.9586 | 1.8924 | 1.72 | 0.086 |  |

## Acknowledgements

The authors thank Prof. Brian Lazzaro (Cornell University, USA) for providing the *Enterococcus faecalis* isolate used in the experiments.

## Author contributions

Conceptualization & Methodology, AB; Investigation, AB and AS; Data curation, Formal analysis, Visualization & Writing (first draft), AB; Writing (review and editing), AB, AS, and NGP; Funding acquisition and Supervision, NGP.

